# Integrative Ensemble Modeling reveals RNA conformations targetable by small molecules

**DOI:** 10.64898/2026.07.28.741152

**Authors:** Stefano Bosio, Vincent Schnapka, Mattia Bernetti, Massimiliano Bonomi, Matteo Masetti

## Abstract

RNA molecules explore heterogeneous conformational ensembles that are essential for their biological function and molecular recognition, yet this intrinsic flexibility poses a major challenge for structure-based drug discovery. In particular, the absence of well-defined binding pockets in static structures limits the identification of ligandable sites. Here, we present an integrative ensemble-based approach that combines enhanced-sampling molecular dynamics simulations with Nuclear Magnetic Resonance data to characterize the conformational landscape of the HIV-1 TAR RNA at atomic resolution. Starting from extensive sampling, we refined the resulting conformational distribution through maximum-entropy reweighting to achieve quantitative agreement with experimental data. Analysis of the reweighted ensemble reveals a diverse set of conformational substates, including compact arrangements that exhibit pocket features compatible with ligand recognition and overlap with known ligand-bound structures. At the same time, highly ligandable conformations, which are only marginally populated, might nonetheless be critical for RNA recognition. Our results demonstrate that integrative ensemble modeling can reveal pharmacologically relevant RNA conformations that are not apparent from experimental static structures, providing a framework for ensemble-based strategies in RNA-targeted drug discovery.

## Introduction

Structure-based drug discovery (SBDD) has become a central paradigm in modern medicinal chemistry.^1^ In its classical formulation, structural information on a macromolecular target is used to rationalize molecular recognition and to guide ligand design to improved potency, selectivity, and physicochemical properties.^1^ This framework has been particularly successful for rigid or moderately flexible proteins, for which a small number of experimentally resolved structures can often be assumed to represent the biologically relevant states. However, this static picture is increasingly recognized as insufficient for targets characterized by pronounced conformational heterogeneity. Proteins with flexible loops, multi-domain architectures, or intrinsically disordered regions, populate broad ensembles of interconverting conformations rather than a single dominant structure.^1–5^ In such systems, ligand recognition frequently involves the selection and stabilization of poorly populated states or, in some cases, substantial ligand-induced rearrangments.^6,7^ A similar principle applies to functional RNAs, whose role as pharmaceutical targets is increasingly recognized.^8–12^ From a computational perspective, most structure-based tools have been historically developed for and benchmarked on protein targets; only the recent surge of interest in RNA molecules as possibly druggable targets has highlighted limitations in this context, motivating the development of RNA-specific and ensemble-based approaches.^4,13–23^

Owing to its intrinsic flexibility resulting from shallow free-energy landscapes, RNA exhibits large motions spanning multiple length and time scales.^24–26^ This dynamics is not merely incidental but directly encodes RNA function, biological recognition, and ligandability.^27,28^ Due to these characteristics, structure-based drug discovery critically depends on an accurate description of RNA conformational ensembles rather than on isolated structural snapshots.^4,14,24,25,29,30^ The HIV-1 trans-activation response element (TAR) is a paradigmatic example of a highly dynamic regulatory RNA and a long-standing target for antiviral intervention.^31^ TAR is a ∼59-nucleotide stem–bulge–loop RNA located at the 5′ end of all nascent HIV-1 transcripts, where it plays a central role in viral replication by recruiting the Tat protein and associated host factors to promote efficient transcriptional elongation.^31–34^ Although TAR is an RNA element of relatively small size, it displays pronounced inter-helical motions, bulge plasticity, and loop rearrangements, which collectively generate a diverse ensemble of transient conformations.^35,36^ Over the years, multiple small molecules, peptides, and peptidomimetics have been shown to bind TAR at distinct sites, including the bulge, the major and minor grooves, and the apical loop (**Figure 1**).^37–40^ Despite decades of structural and biochemical investigations, the rational discovery of small-molecule modulators of HIV-1 TAR remains challenging. This partly reflects the absence of a single, well-defined binding pocket in unbound TAR, which instead populates a heterogeneous ensemble of transient conformations.^41,42^ The dynamic nature of HIV-1 TAR presents both an opportunity and a challenge for SBDD. On the one hand, conformational flexibility can give rise to transient pockets and alternative interaction surfaces that are not apparent in any single static structure, thereby expanding the landscape of exploitable binding sites. On the other hand, the sparsity and low population of many of these substates complicate their direct experimental characterization and hinder the selection of appropriate structural representatives for computational screening.

**Figure 1.**
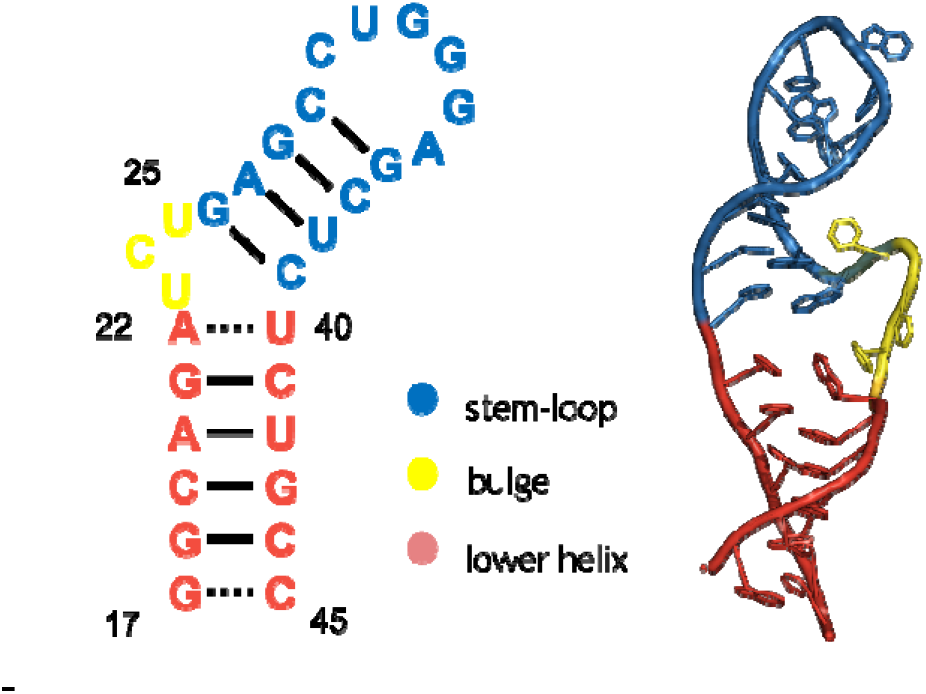
Structural representation of the HIV-1 TAR RNA region investigated in this study. Secondary (left) and three-dimensional (right) representations of HIV-1 TAR RNA (PDB ID: 1ANR^45^) highlighting the nucleotide segment included in the simulations (nucleotides 17–45). The apical loop, bulge, and lower helical stem are shown in distinct colors to emphasize the structural elements that dominate TAR conformational dynamics and the recognition from the biological partners.

More generally, the growing recognition that molecular function emerges from dynamic ensembles has underscored the need for integrative structural biology approaches that couple physics-based simulations with quantitative experimental information. Such frameworks provide a mechanistic bridge between atomistic conformational populations and functional readouts.^43,44^ These considerations motivate the development of strategies that combine extensive conformational sampling with experimental restraints, enabling a quantitative, ensemble-level description while preserving atomistic resolution.

In this study, we combine enhanced-sampling molecular dynamics (MD) simulations with Nuclear Magnetic Resonance (NMR) residual dipolar couplings (RDCs) to generate an ensemble of HIV-1 TAR consistent with experimental data (**Figure 2**). We further validate the resulting ensemble against multiple independent experimental measurements. We then characterize the conformational substates, examine their relationship to known ligand-bound structures, and identify those exhibiting pocket amenable to structure-based drug design. Our results show that reweighting guided by experimental data reshapes the HIV-1 TAR conformational ensemble while preserving its structural heterogeneity. These findings highlight how integrative ensemble modeling can uncover pharmacologically relevant RNA conformations that are not apparent from static structures alone, opening new avenues for RNA-targeted drug discovery.

**Figure 2.**
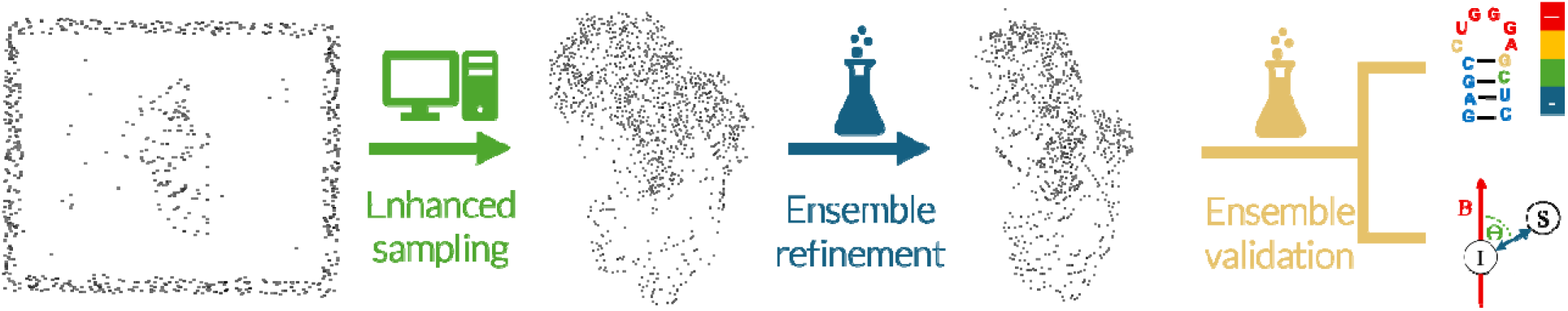
Integrative workflow for ensemble-based characterization of HIV-1 TAR RNA. Enhanced-sampling MD simulations are used to extensively explore the conformational dynamics of HIV-1 TAR RNA and generate an initial ensemble of structures, capturing both large-scale inter-helical motions and local structural fluctuations. This prior ensemble is subsequently refined using NMR experimental data through maximum-entropy reweighting, to obtain an experimentally-consistent distribution while preserving conformational heterogeneity. Finally, the refined ensemble is validated against independent experimental data, including RDCs and SHAPE reactivity.

## Methods

### Simulations setup and protocol

The initial atomic coordinates were extracted from the NMR structure of the HIV-1 TAR RNA (PDB ID: 2KX5^30^). RNA parameters were assigned using the AMBER99^46,47^ force field with the BSC0 correction on torsional angles^48^ and the correction on anti-g shift (χOL3) ^49^ as implemented in OpenMM library v 7.7.0.^50^ The Joung–Cheatham ion parameters^51^ and the OPC water model^52^ were employed. The system was solvated in a truncated octahedral box of explicit water molecules, ensuring a minimum distance of 20 Å between any RNA atom and the box boundary to allow for large-scale conformational fluctuations. Potassium and chloride ions were added to neutralize the system and to reproduce a bulk salt concentration of 0.15 M. Electrostatic interactions were treated using the smooth particle mesh Ewald method with a cutoff of 9 Å.^53^ Van der Waals interactions were smoothly switched off between 8 Å and 9 Å. The equations of motion were integrated with the leap-frog algorithm using a 2 fs time step. Short parallel tempering (PT)^54^ simulations were performed using 24 replicas geometrically distributed between 300 K and 450 K, equilibrating each one at its target temperature through a sequential heating protocol. The system was equilibrated through a sequential heating protocol, starting from 300 K. Each subsequent replica was initialized from the final configuration of the previous one and simulated for 1 ns at its target temperature. A Parallel Tempering in the Well-Tempered Ensemble (PT-WTE)^55^ simulation was then conducted for a total of 16.8 μs, using the system potential energy as the collective variable. Gaussians with height *W* = 2.4 kJ mol ¹ and width σ = 500 kJ mol ¹ were deposited every 500 steps. A bias factor of 25 was applied to ensure sufficient overlap between the energy distributions of neighboring replicas, and replica exchanges were attempted every 100 steps, yielding an average acceptance rate of 45.6%. All MD simulations were carried out using GROMACS 2023.2^56–58^ equipped with PLUMED 2.9.^59–61^

### Calculation of experimental observables and reweighting strategy

Ensemble-averaged RDCs were obtained by calculating the couplings for each MD snapshot using PALES.^62,63^ The calculations were performed within a cylindrical wall potential, employing a *pf2* effective concentration of 0.022 g mL□¹ and assuming a rod-like liquid crystal model in agreement with the experimental conditions of the reference values.^64,65^ The alignment tensor was determined globally across the ensemble. To account for differences in alignment magnitude between simulations and experiments, the predicted RDCs were multiplied by one global scaling factor per dataset.

SHAPE reactivity is known to correlate with the local flexibility of nucleotides.^66,67^ In this study, we followed established approaches to investigate the relationship between SHAPE reactivity and flexibility parameters derived from MD simulations.^66^ We compared experimental SHAPE reactivity profiles with an MD-derived flexibility metric based on inter-nucleotide distance fluctuations between consecutive residues, with the per-residue value obtained by averaging the fluctuations of the adjacent pairs [*i*–1,*i*] and [*i*,*i*+1] as a measure of the local flexibility of residue *i*. The experimental SHAPE reactivity profiles used in this analysis were extracted from previously published measurements.^68^

Maximum-entropy (MaxEnt) reweighting provides a theoretical framework to integrate experimental data into MD simulations while minimally perturbing the original prior ensemble.^69,70^ By enforcing agreement with solution experimental observables while preserving ensemble heterogeneity, this approach is particularly well suited for flexible biological systems.^71,72^ The experimental data were enforced following the formulation described in Cesari et al. through custom in-house scripts.^70,73^ The procedure determines a set of Lagrange multipliers λ*ᵢ* so that ensemble-averages are computed with weights *W_t_* matching reference experimental values:

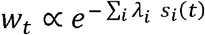

Where *s_i_(t)* is the value of the *i*th observable computed for structure *t* of the prior ensemble; *λ_i_* denotes the optimal Lagrange multipliers that minimize the following quantity:

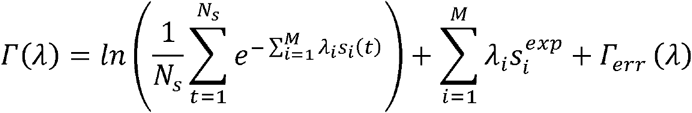

Where *N_s_* is number of structures in the prior ensemble, *s^exp^_i_* is the experimental value of the ith observable, and *Γ_err_ (λ)* is an error term that accounts for the uncertainty associated with the experimental measurements. The latter acts as a regularization term to prevent overfitting and is defined as:

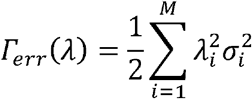

Where *σᵢ* specifies the uncertainty associated with the observable *Sᵢ*. The subset of prior configurations contributing significantly to the reweighted ensemble can be estimated via the Kish effective sample size:

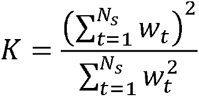

Herein, MaxEnt reweighting was performed using previously published RDC measurements,^64,74^ and the resulting agreement with the experimental data evaluated via the Pearson correlation.

## Results

### Maximum entropy reweighting yields an HIV-1 TAR ensemble consistent with experiments

The conformational space of HIV-1 TAR was thoroughly explored using the PT-WTE enhanced-sampling approach. We then applied MaxEnt reweighting to modulate the statistical weights of the conformations from the PT-WTE prior ensemble so that ensemble-averaged observables match experimental data within their uncertainties. RDCs were chosen as target observables, as they report on average internuclear vector orientations relative to the magnetic field and thus provide long-range information that are highly sensitive to inter-helical motions and relative domain orientations. This refinement preserves ensemble diversity and keeps the reweighted distribution close to the PT-WTE prior, mitigating overfitting through the optimization of the regularization strength (σ). To determine the optimal σ, we performed 10-fold cross-validation with randomized splits: in each fold, 90% of RDCs were used to fit the MaxEnt multipliers and the remaining 10% were held out to compute the root-mean-square error (RMSE) (**Figure 3E**). For each σ, we averaged the RMSE across folds to obtain a robust estimate of predictive performance and selected the value which minimizes this metric. Using the selected σ (7 Hz), we recalculated the weights and recomputed the RMSE over all observables. We also monitored the RMSE on the test set and the Kish effective sample size to diagnose potential overfitting. Back-calculated RDCs from the prior PT-WTE ensemble (**Figure 3A** green) deviate systematically from experiments (**Figure 3A** grey), with larger discrepancies in the apical loop and bulge. Quantitatively, the initial agreement was modest (R≈0.68, RMSE = 8.4 Hz).

**Figure 3.**
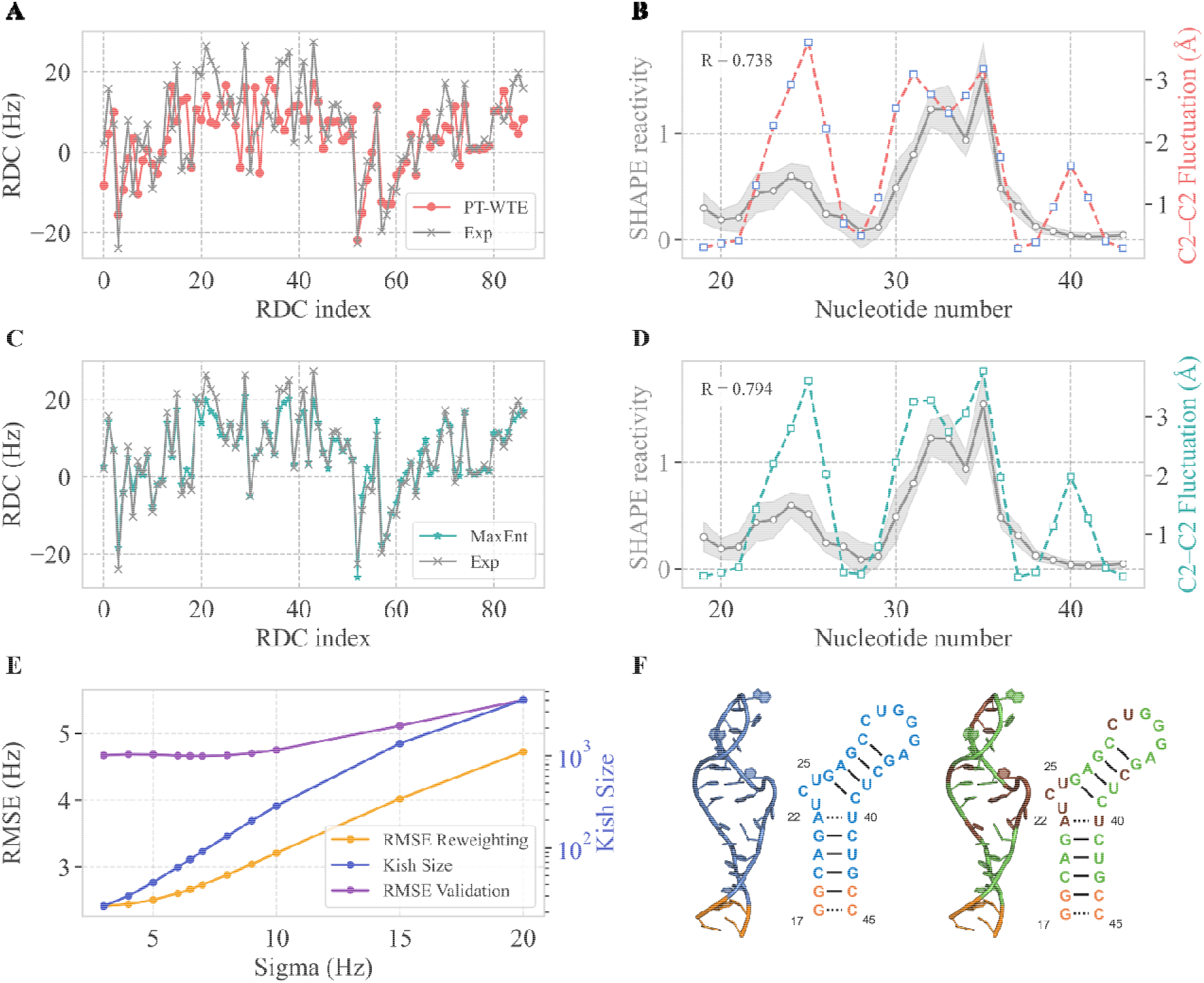
Maximum entropy reweighting of TAR structural ensembles against RDCs and validation with independent data. (A) Comparison between experimental RDC values (grey) from previous NMR studies^74,75^ and back-calculated RDCs from the prior PT-WTE ensemble (green). (B) Comparison between experimental SHAPE reactivity profiles (grey) and computed inter-nucleotide distance fluctuations between consecutive residues, with the per-residue metric obtained by averaging the fluctuations of adjacent pairs. (C) Comparison between experimental RDC values (grey) from previous NMR studies and back-calculated RDCs from reweighted back-calculated RDCs (red). (D) Comparison between experimental SHAPE reactivity profiles (grey) and computed fluctuations in the mean inter-nucleotide distances between consecutive residues after reweighting (red). (E) Cross-validation scheme for optimization of the σ parameter. The root means square error (RMSE) obtained from 10-fold validation of the RDC dataset (violet) is plotted as a function of σ. The associated Kish size is reported in blue, whereas the RMSE computed over the full dataset is shown in yellow. (F) Structural representation of TAR highlighting the residues included (blue) or excluded (orange) from SHAPE-based validation (right); residues correctly reproduced by the reweighted ensemble are shown in green, while those with the largest deviations are marked in brown(right).

The cross-validated selection of σ yielded a reweighted ensemble with improved agreement with held-out RDCs while preserving a substantial effective sample size. After maximum-entropy reweighting, the agreement increased significantly (Figure 3C): the correlation with experimental RDCs increased to R ≈ 0.97, while the RMSE decreased to 2.8 Hz. Cross-validated test errors displayed a minimum at intermediate σ values, indicating that the selected regularization balances fitting accuracy. While agreement with the experimental data improved, the Kish effective sample size (K = 74.2 over a total number of 70,000 structures) indicated that the reweighted ensemble concentrates weight on a relatively small subset of structures.

As independent validation, we compared the reweighted ensemble against SHAPE reactivity data from *ex virio* and transcript samples (Figure 3B-D, Figure S1), neither of which were used for refining. In both cases, the reweighted ensemble reproduced the experimental profiles more accurately than the prior PT-WTE ensemble. Notably, the correlation with SHAPE reactivities in the transcripts data slightly increased from 0.73 to 0.79 upon reweighting and a similar trend is observed in the *ex virio* data.

### Reweighted ensemble reveals HIV-1 TAR structural heterogeneity

To characterize the conformational space explored in our simulations we computed G-vectors,^76^ a feature set suited to RNA structures that is sensitive to subtle rearrangements of base pairing/stacking that can nucleate or disrupt ligandable pockets. As a reference, we collected experimental HIV-1 TAR structures from the PDB in the unbound (red) and ligand-bound (blue) states and projected them onto the same reduced space. The representation given by dimensionality reduction through PCA on G-vectors allows us to visualize the high-dimensional manifold in two dimensions. The projection (**Figure 4A** right) shows that the prior PT-WTE ensemble spans a broad range of conformations that envelop the regions populated by both unbound and ligand-bound PDB structures.

**Figure 4.**
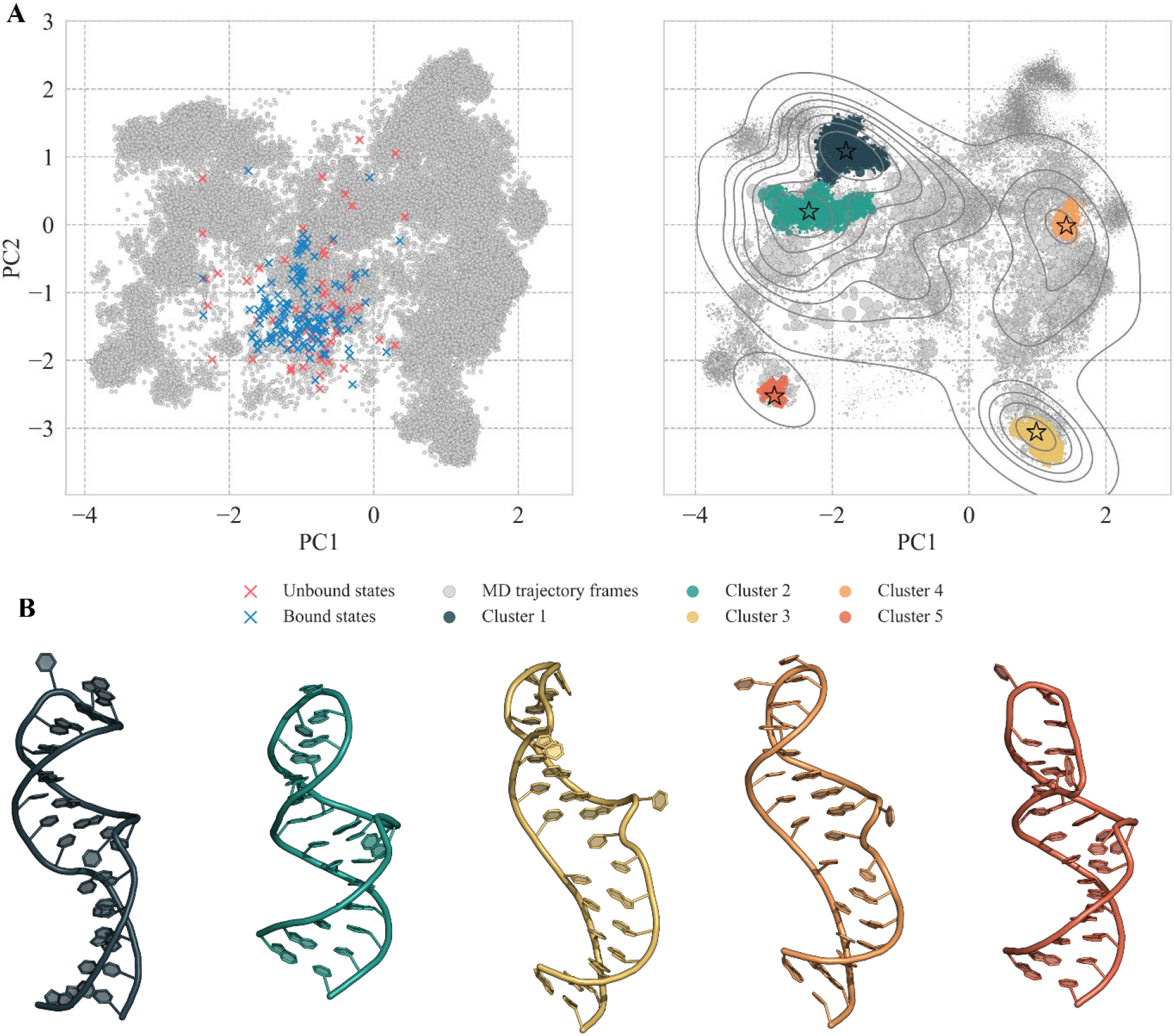
Conformational space of TAR before and after maximum entropy reweighting. (A) Two-dimensional projection of the conformational space sampled during the PT-WTE simulation (grey points), compared with reference PDB structures of TAR in the unbound (red) and bound (blue) states. The left panel shows the points distribution before reweighting, while the right panel displays the same projection after applying MaxEnt reweighting. Grey contour lines trace isodensity levels from a kernel density estimate computed with the MaxEnt weights, highlighting high probability basins and their spatial extent. Eight levels are shown with a bandwidth adjustment of 0.65 and a density threshold of 0.02. PC1 and PC2 explain 16.72% and 11.25% of the total variance, respectively. Representative structures of the main clusters are indicated by star markers. (B) Representative structures of the main clusters identified in the reweighted ensemble, colored according to the scheme used in panel A.

After maximum-entropy reweighting (**Figure 4A**, left), populations shift toward unbound-like states, including regions not sampled by available PDB structures. At the same time, overly extended conformations that dominated the prior are de-emphasized, as in the case of cluster 3 (**Figure 4B** yellow) and cluster 4 (**Figure 4B** orange), and few discrete conformations that overlap with ligand-bound crystallographic references gain statistical relevance. These shifts in the ensemble are reflected in the representative post-reweighting conformations, which reveal a range of inter-helical arrangements (**Figure 4B**). Cluster 1 is compact and near-coaxial, with a modestly splayed bulge and the apical loop folded toward the stem. Cluster 2 remains compact but exhibits a mild inter-helical twist that extrudes the bulge and opens a narrow cleft along the minor groove. Conversely, clusters 3 and 4 are more extended, with the helical arms splayed and the loop projected outward, increasing solvent exposure but yielding a diffuse, poorly defined cavity. Finally, cluster 5 is compact yet asymmetric, with the loop bending toward the bulge to form transient tertiary contacts that may stabilize a lateral microcavity or modulate access to the principal pocket.

We next determined ligandability scores through the SiteMap software^77,78^ and mapped them onto the PCA landscape to relate geometry with pocket quality (**Figure 5** and **Figure S2**). In the WTE prior, scores span a broad range, with a dense cloud of low SiteScore corresponding to overextended conformations that lack well-defined hotspots for small-molecule recognition. Outside this region, most regions exhibit medium–to–high SiteScore values, typically associated with compact or kinked inter-helical arrangements. Interestingly, after MaxEnt reweighting the probability of high-ligandable states decreases markedly, with population shifting toward medium–low ligandable basins. Nonetheless, a small subpopulation of elongated conformations persists with non-negligible statistical weights.

**Figure 5.**
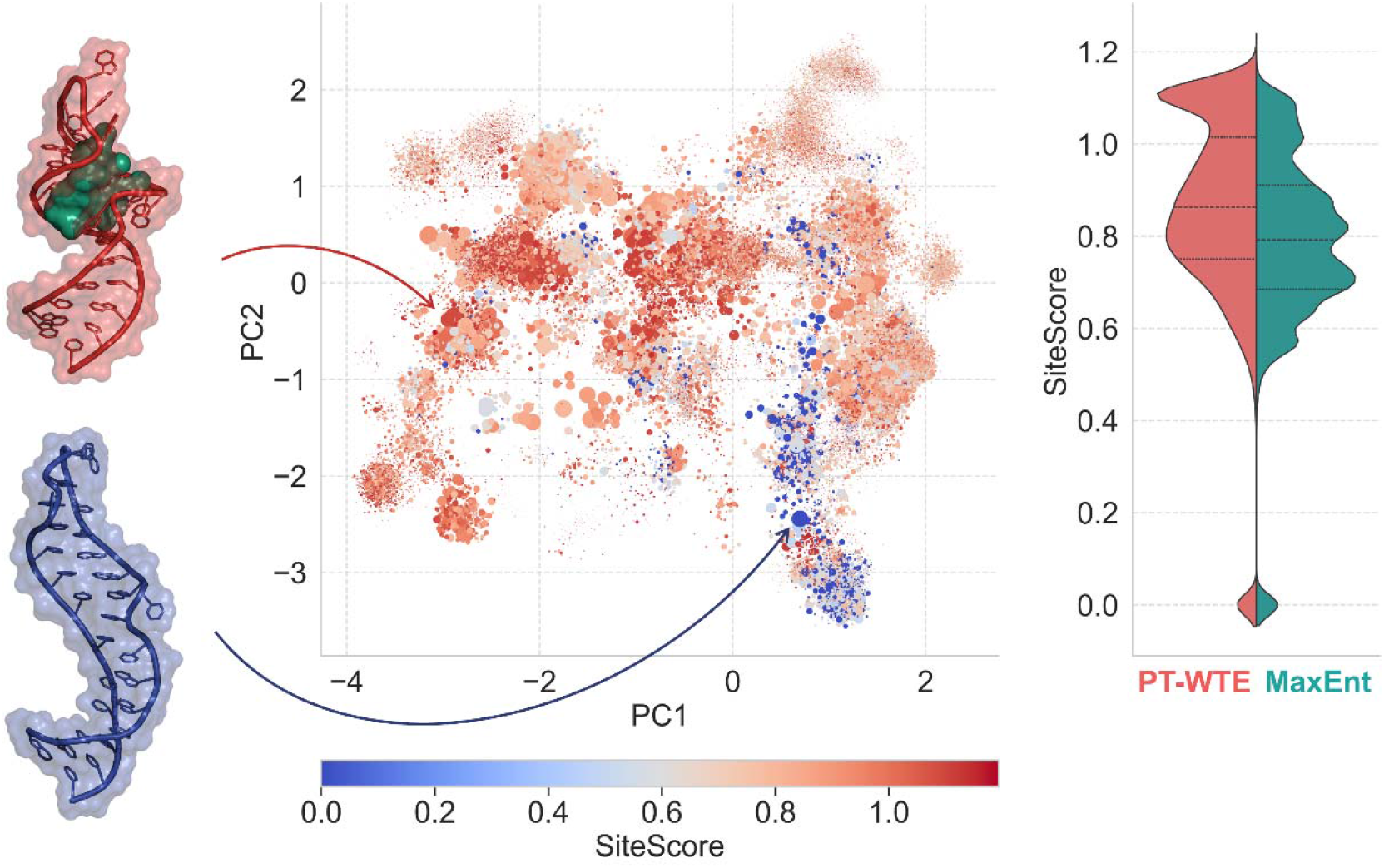
SiteScore across the HIV-1 TAR conformational space. The central panel shows the projection onto the first two principal components (PC1 and PC2), where each point represents a conformational state. Colors encode the predicted SiteScore values, where lower scores indicate less ligandable conformations and higher scores indicate more ligandable ones. The color scale rangesfrom blue (low ligandability) to red (ligandability), while point sizes are scaled according to the statistical weights of the conformations. Two representative regions are highlighted: a high-scoring basin (red), where the best-scoring conformation is shown together with its associated binding pocket (green), and a low-scoring region (blue), corresponding to the worst-scoring conformation. The right panel reports the SiteScore distributions accounting for MaxEnt reweighting, comparing the original ensemble (red) with the reweighted ensemble (green).

### HIV-1 TAR exhibits partial pre-organization for ligand-bound conformations

To quantify the relative populations of conformers resembling known TAR-ligand complexes, we first examined the region of conformational space proximal to the available experimental structures. To this end, we performed cluster analysis via the QT algorithm^79^ using the same metrics adopted for dimensionality reduction. We then evaluated cluster populations as a function of increasing εRMSD^76^ from the respective references and identified the corresponding cluster representatives (**Figure S3**).

Comparison between the prior and reweighted ensembles shows that the 1QD3 structure^38^ remains consistently represented across all εRMSD thresholds, with only minor changes in statistical weight. At the most stringent cutoff, approximately 5% of the ensemble populates conformations whose geometry is compatible with the 1QD3 binding pocket. A similar trend is observed for 1UTS,^40^ although populations below an εRMSD of 0.9 are limited. At higher thresholds, the cumulative weight associated with 1UTS decreases after reweighting. Increasing the cutoff to 0.9 brings the 1UUI^80^ and 2L8H^81^ references into the ensemble, with 2L8H showing a marked increase in population after reweighting. At a cutoff of 1.0, 1UUD^80^ also become populated, whereas 1LVJ^82^ remains poorly represented, with the closest cluster lying beyond the range explored.

A further analysis of the pocket rearrangement was performed by assigning each reference structure to the closest cluster within 5 kcal mol^-1^ of the global minimum, and projecting the experimentally known pocket conformations onto the principal-component space derived from the PT-WTE ensemble (**Figure S4**).. Among the six experimental cases examined, three structures (1QD3, 2L8H, and 1UUD) display substantial overlaps with regions of high probability density in the simulated ensemble, therefore showing a high degree of pre-organization. For 1UTS, the crystallographic conformations fall in regions of lower density. By contrast, 1LVJ and 1UUI exhibit limited overlap with the sampled conformational space. An illustrative example is provided by 1QD3 (**Figure 6A**). The nearest unbound representative lies within the same basin as the crystal structure in the PCA space of the pockets, with a crystal– centroid εRMSD of 0.72. Similar geometric agreement is observed for 1UUD and 1UUI **(Figure S5** and **Figure S6**), where the ligands occupy solvent-exposed regions, and the interacting surfaces remain largely preserved. For 1UTS, the closest representative is found at an εRMSD of 0.77. In this case, the ligand binds in the bulge region, and repositioning of nucleotides U23 and U40 is required to accommodate the bound state (**Figure S5**). The G10–C23 base pair is maintained but undergoes a modest reorientation. The 2L8H complex involves a pocket located within the apical loop. In the closest unbound representative, G36 partially occludes the cavity and must shift by approximately 3.5 Å to allow ligand accommodation. A35 is observed in an extrahelical conformation, while G34 transitions between extrahelical and intrahelical states to stabilize the bound geometry (Figure S6). Finally, for 1LVJ, the nearest non-negligible representative is found at an εRMSD of 1.11 from the crystallographic structure. In this case, the G26–C13 base pair is retained, but its relative orientation would sterically clash with the ligand. Additional rearrangements involve loss of stacking between C24 and U25 and disruption of the U23–U40 base pair to permit ligand accommodation (**Figure 6B**).

**Figure 6.**
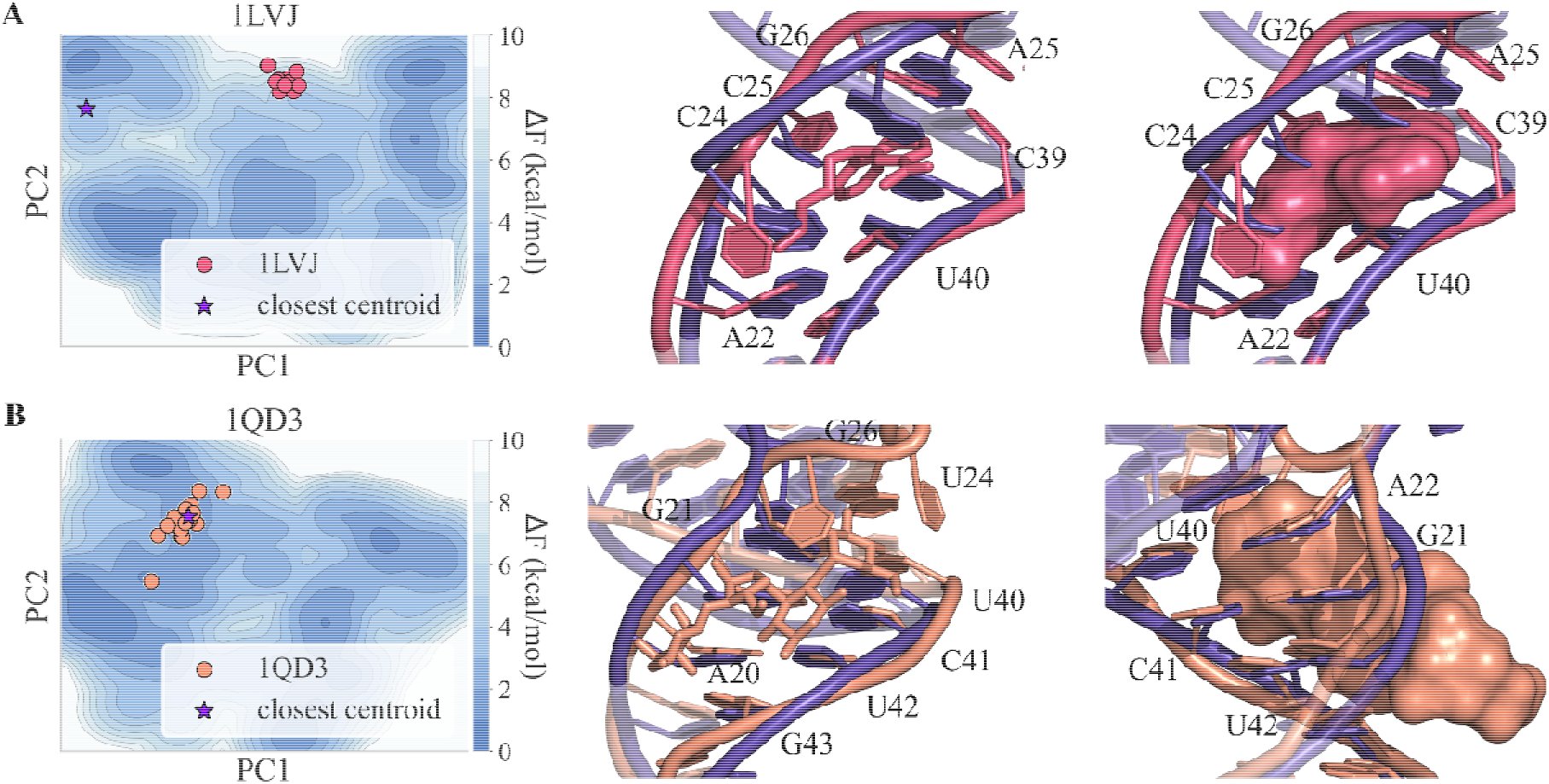
Structural pre-organization of ligand-bound TAR conformations within the unbound ensemble. (A) Two-dimensional projection of the conformational free-energy landscape sampled around the binding mode observed in the 1LVJ structure, shown along the first two principal components (PC1 and PC2). Colored circles correspond to the crystallographic poses of the ligand in 1LVJ, and the purple star indicates the centroid of the closest simulated cluster. In the center, the three-dimensional comparison between the crystallographic binding mode and the closest simulated conformation is shown. The ligand is represented in stick form, and HIV-1 TAR is displayed as a cartoon and stick representation. Residue numbers are indicated. On the right, the same view and color scheme are used, but the ligand is represented as a molecular surface to illustrate the occupied binding cavity. (B) Two-dimensional projection of the conformational free-energy landscape sampled around the binding mode observed in the 1QD3 structure, shown along PC1 and PC2. Colored circles correspond to the crystallographic poses of the ligand in 1QD3, and the purple star indicates the centroid of the closest simulated cluster. In the center, the three-dimensional comparison between the crystallographic binding mode and the closest simulated conformation is shown. The ligand is displayed in stick representation from the front view, and HIV-1 TAR is shown as a cartoon and stick representation. On the right, the ligand is represented as a molecular surface after a 180° rotation to highlight shape complementarity.

## Discussion

A major challenge in describing highly dynamic RNAs such as HIV-1 TAR is reconciling extensive conformational sampling with quantitative agreement to experimental data, while retaining a physically meaningful ensemble representation. Recent works have increasingly framed RNA structure determination as an ensemble problem, rather than the identification of a single representative conformation, particularly for TAR, where low-populated substates have been shown to play a function role in recognition and regulation.^83^

Maximum entropy reweighting against RDCs substantially improves agreement with experimental data while avoiding an artificial collapse of the ensemble onto an extremely low number of conformations. The observed redistribution of statistical weights across substantially different states sampled during PT-WTE simulations highlights the potential of enhanced sampling approaches in capturing the heterogeneity/complexity of TAR conformational landscape. On the other hand, the limited Kish effective sample size observed after reweighting reflects the well-known difficulty of modeling extremely flexible biomolecules..

Independent validation using SHAPE reactivity data further supports the internal consistency of the refined ensemble. Unlike RDCs, which are well-defined ensemble-averaged observables that can be computed unambiguously for each conformation, SHAPE reactivities report indirectly on local flexibility, base-stacking propensity, and solvent accessibility.^66^ As a result, as of today no rigorous structural forward model exists to predict SHAPE values directly from individual conformations, preventing their inclusion as a target in the maximum entropy functional. Nevertheless, the refined ensemble shows a modest but systematic improvement in agreement with SHAPE profiles that were not used during fitting, suggesting that refinement against observables accounting for long-range interactions can indirectly enhance the description of local structural features. This is consistent with previous findings indicating a strong coupling between global RNA motions and local flexibility patterns.^66^ The deviations from experimental data likely reflect a combination of the aforementioned methodological issues in computing SHAPE reactivity and system-specific factors. In addition, in this specific case, the simulation targets a pharmacologically relevant TAR region, whereas SHAPE measurements were obtained for the full-length transcript, where long-range interactions and global architecture may influence local flexibility patterns beyond the scope of the truncated model.

Analysis of the conformational landscape before and after reweighting indicates that maximum entropy reshapes the population distribution while largely preserving the structural heterogeneity of the original ensemble. Notably, only a limited number of microstates are substantially de-emphasized upon reweighting, whereas isolated conformations in the vicinity of experimentally resolved crystal structures gain statistical weight. These changes highlight the importance of extensive prior sampling and an accurate force field capable of describing physiologically relevant conformations: experimental refinement can only act on substates that are already explored, but it enables a clearer separation between conformations consistent with experimental observables and those that are not.

Mapping pocket metrics onto the conformational landscape reveals that ligandability is a population-weighted ensemble property rather than an intrinsic attribute of any single structure. Compact and mildly kinked inter-helical conformations tend to support more defined pockets, whereas highly extended states lack persistent features suitable for small-molecule recognition. After reweighting, the probability of highly ligandable conformations decreases, indicating that such states could be intrinsically rare in the unbound ensemble. This observation underscores a fundamental challenge in RNA-targeted drug discovery: ligand-accessible pockets may correspond to low-population substates that nevertheless become functionally relevant upon ligand binding.

In this context, it is important to emphasize that, particularly for RNA targets, pocket-based druggability scores such as those reported by SiteMap should be interpreted as descriptive metrics rather than quantitative predictors of binding.^84^ Unlike many protein targets, RNA molecules can engage small molecules through transient, shallow, or highly dynamic interaction surfaces that may not correspond to well-defined pockets in a static structural sense. Consistent with this view, several RNA-targeting ligands have been identified despite the absence of a canonical binding pocket, as exemplified by the recently discovered drug Risdiplam.^85^ In such cases, the extreme conformational flexibility of the target places RNA in close conceptual analogy with intrinsically disordered proteins. In these systems, ligand recognition can occur through the stabilization of specific conformational subensembles, driven primarily by entropic contributions rather than the formation of a single, well-defined and enthalpy-driven binding mode.86,87

A systematic comparison between the refined ensemble and available ligand-bound TAR structures reveals that ligand recognition spans a continuum between conformational selection and induced fit, rather than conforming to a strict dichotomy.^88^ Several complexes, such as those represented by the PBD structures 1QD3, 2L8H, and 1UUD, are compatible with pre-organized conformations explored by the unbound ensemble, differing only by modest local rearrangements. In contrast, other complexes, most notably 1LVJ, require more extensive reorganization of base pairing and stacking interactions that are rarely visited in the unbound ensemble. Intermediate cases, such as 1UTS and 1UUI, occupy regions of low but non-negligible probability density, suggesting that their corresponding pocket geometries are accessible yet underrepresented prior to ligand binding. Together, these observations support a continuum model of TAR recognition, in which the balance between conformational selection and ligand-induced stabilization depends on the binding site and the chemical nature of the ligand.^88^

From a structure-based drug discovery perspective, these results highlight the limitations of static representations when targeting highly dynamic RNAs. Small sets of representative structures are unlikely to capture the diversity of conformations relevant for binding. Experimentally refined conformational ensembles instead provide the natural framework for prioritizing substates in docking and ligand optimization workflows. In the case of TAR, compact and mildly twisted conformations emerge as particularly relevant targets, whereas highly extended conformations represent a more challenging target for conventional medicinal chemistry strategies. More broadly, the workflow presented here offers a general route to identify experimentally supported RNA substates and to rationalize ligand recognition mechanisms in terms of population shifts within dynamic ensembles.

Despite these advances, limitations remain. The effective sample size after reweighting indicates incomplete sampling of slow collective motions and rare local rearrangements. Addressing these limitations will require diverse sampling strategies to generate initial ensembles, diverse starting structures, and incorporation of orthogonal experimental restraints, such as from SAXS or NOEs data. Extending this framework to other regulatory RNAs will be essential to assess its generalizability and to further elucidate how RNA dynamics shape ligandability across diverse systems.

## Conclusions

In this work, we present an integrative ensemble-based framework to explore targetable conformations in highly dynamic RNA systems, using HIV-1 TAR as a paradigmatic case study. By combining extensive enhanced-sampling MD with maximum entropy reweighting against NMR RDCs, we obtained an experimentally consistent conformational ensemble that preserves the intrinsic heterogeneity of TAR while enabling a quantitative analysis of its ligand accessible substates. This approach provides a route to move beyond static structural representations in computational drug discovery, particularly for RNA targets. Our results suggest the coexistence of multiple populations, each of them characterized by a unique rearrangement of base pairing and stacking.

Our work emphasizes the need for ensemble-aware strategies in RNA-target drug discovery, in which experimentally refined conformational distributions guide the prioritization of relevant substates for subsequent docking, virtual screening, or ligand optimization. With growing interest in RNA as a therapeutic target, integrative approaches that explicitly capture structural heterogeneity will be essential for rationalizing ligand recognition and extending SBDD to highly dynamic biomolecular systems.

## Data availability

We freely provide the input files (initial coordinates, topologies, GROMACS mdp parameter file, and PLUMED inputs) to perform the MD simulations in this work, as well as the output simulation trajectories (in xtc format) that we generated. We supply a Jupyter notebook to reproduce our analyses, results, and the plots reported in this work. All the material is freely available in Zenodo with accession code 10.5281/zenodo.21352447. PLUMED input files are also available on the PLUMED-NEST under plumID:26009, while the Jupyter notebook can also be straightforwardly consulted and downloaded at github.com/CompMedChemLab/tar-ensemble-modeling. All MD simulations were performed with GROMACS 2023.2, equipped with plumed 2.9.

## Supporting Information

The following files are available free of charge. Comparison of experimental SHAPE reactivity profiles with MD-derived metrics for the ex virio dataset (**Figure S1**); PCA projection of the TAR conformational ensemble sampled by PT-WTE, annotated with SiteScore and with/without MaxEnt statistical weights (**Figure S2**); population of TAR conformations within increasing εRMSD thresholds relative to ligand-bound PDB references, before and after reweighting (**Figure S3**); two-dimensional free-energy landscape of TAR binding-pocket conformations in principal-component space, with ligand-bound pocket geometries and nearest simulated cluster centroids overlaid (**Figure S4**); and three-dimensional structural comparisons between the closest ensemble representatives and corresponding ligand-bound crystal structures (**Figures S5–S6**).

## AUTHOR INFORMATION

Contributions: M. Bonomi and M. Masetti conceived the study. S. Bosio, V. Schnapka, and M. Bernetti performed the molecular dynamics simulations, conducted the investigation, and curated the data. All authors contributed to writing and revising the manuscript and approved the final version.

## ACKNOWLEDGMENT

The authors would like to thank Paraskevi Gkeka for useful discussions.

We gratefully acknowledge the Data Science and Computation Facility and its Support Team for their support and assistance with the IIT High Performance Computing Infrastructure.

