## Supplementary Material for "Integrative Ensemble Modeling reveals RNA conformations targetable by small molecules"

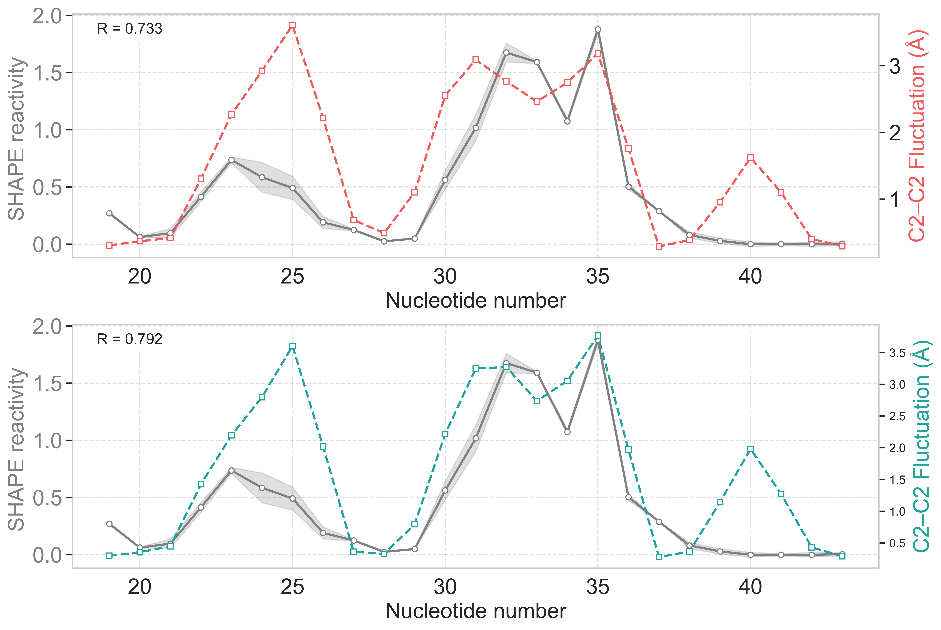

**FIG S1 Comparison of experimental SHAPE reactivity profiles with MD-derived metrics for the ex virio dataset**. Experimental SHAPE reactivity profiles (grey) and computed fluctuations in the mean inter-nucleotide distances between consecutive residues calculated on the *ex virio* data. Pre-reweighting data are shown in red, and reweighted data in green.

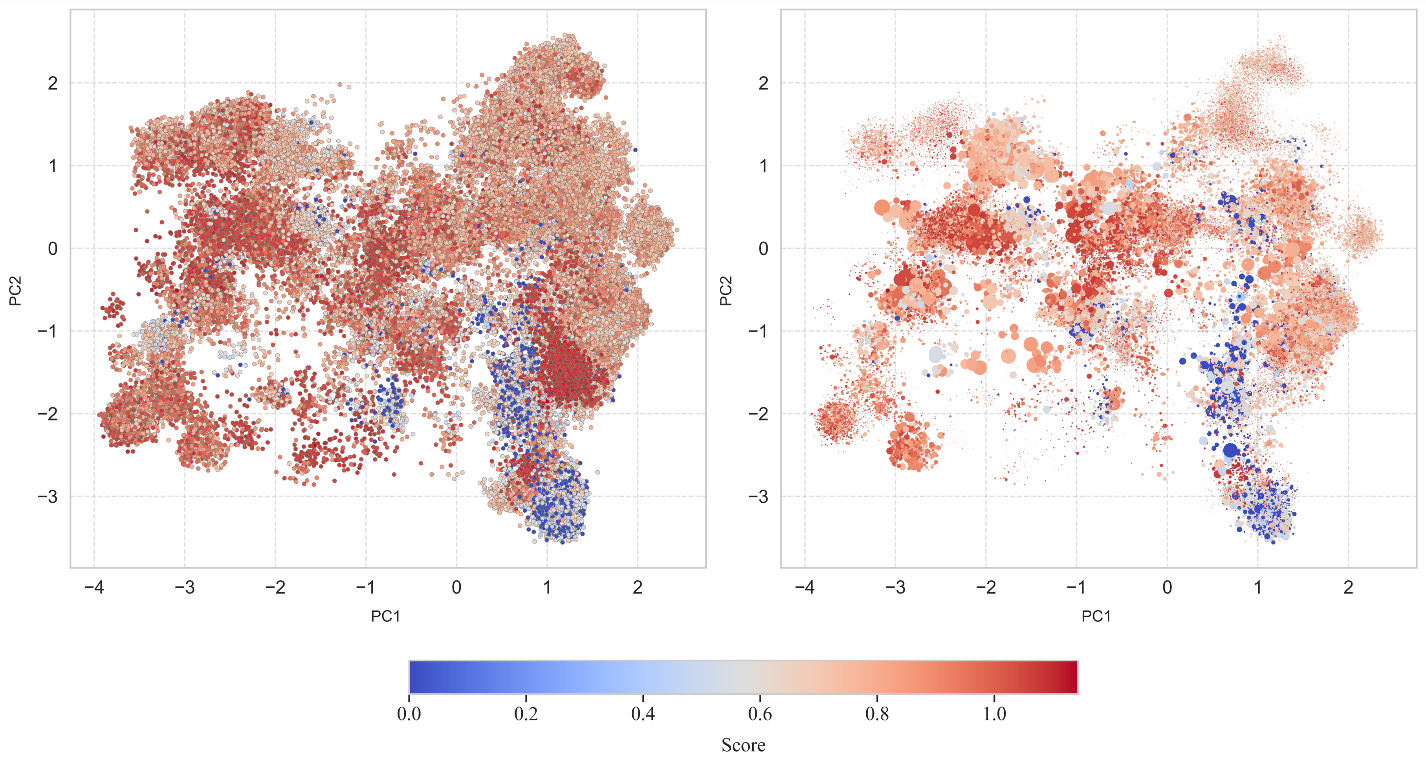
**FIG S2 Two-dimensional projection of the conformational space sampled during the WTE simulation.** Colors indicate SiteScore values, with blue corresponding to low values and red corresponding to high values. On the right, point sizes are proportional to the statistical weight of each conformation. On the left, point sizes are constant because all weights are equal.

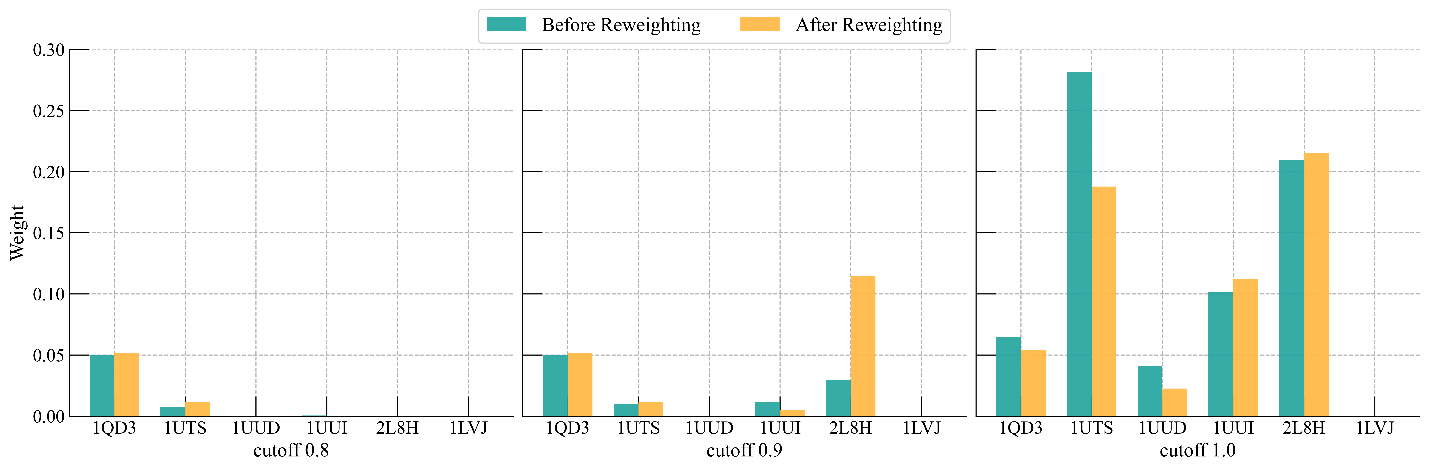

**Figure S3 Population of TAR conformations within increasing εRMSD thresholds relative to ligand-bound PDB references, before and after reweighting**. The green bars correspond to the pre-reweighting ensemble, while the yellow bars represent the reweighted ensemble.

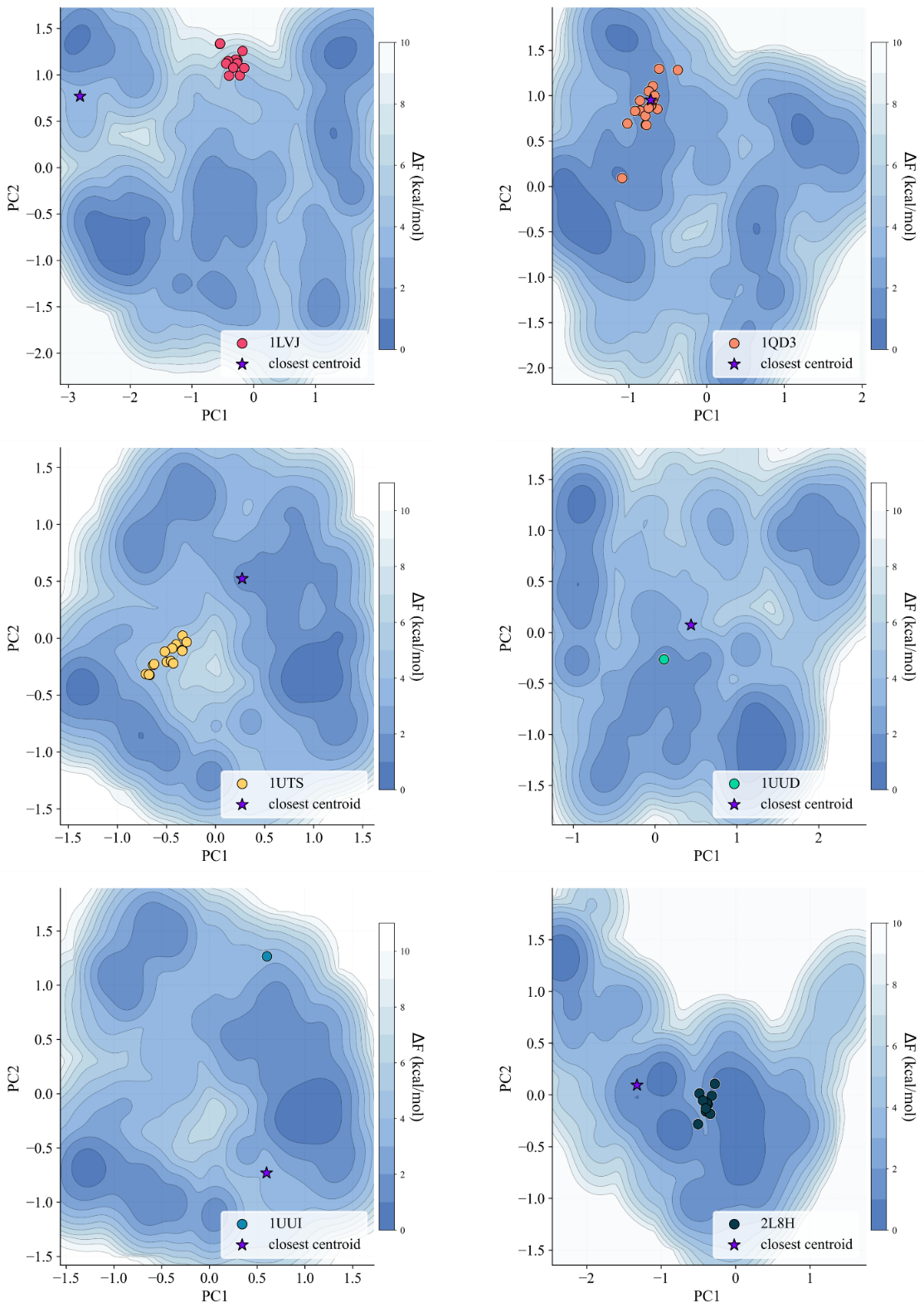

**Figure S4 Two-dimensional projection of the conformational free-energy landscape of TAR binding pockets.** Free-energy surfaces were computed for pocket conformations sampled during the PT-WTE simulations along the first two principal components (PC1 and PC2). Colors indicate the relative free-energy values, from low-energy (dark blue) to high-energy (light blue) regions. Experimentally determined ligand-bound pocket conformations are overlaid as colored dots, with colors corresponding to the respective PDB identifiers. Purple stars indicate the centroids of the simulated clusters closest to each experimental conformation

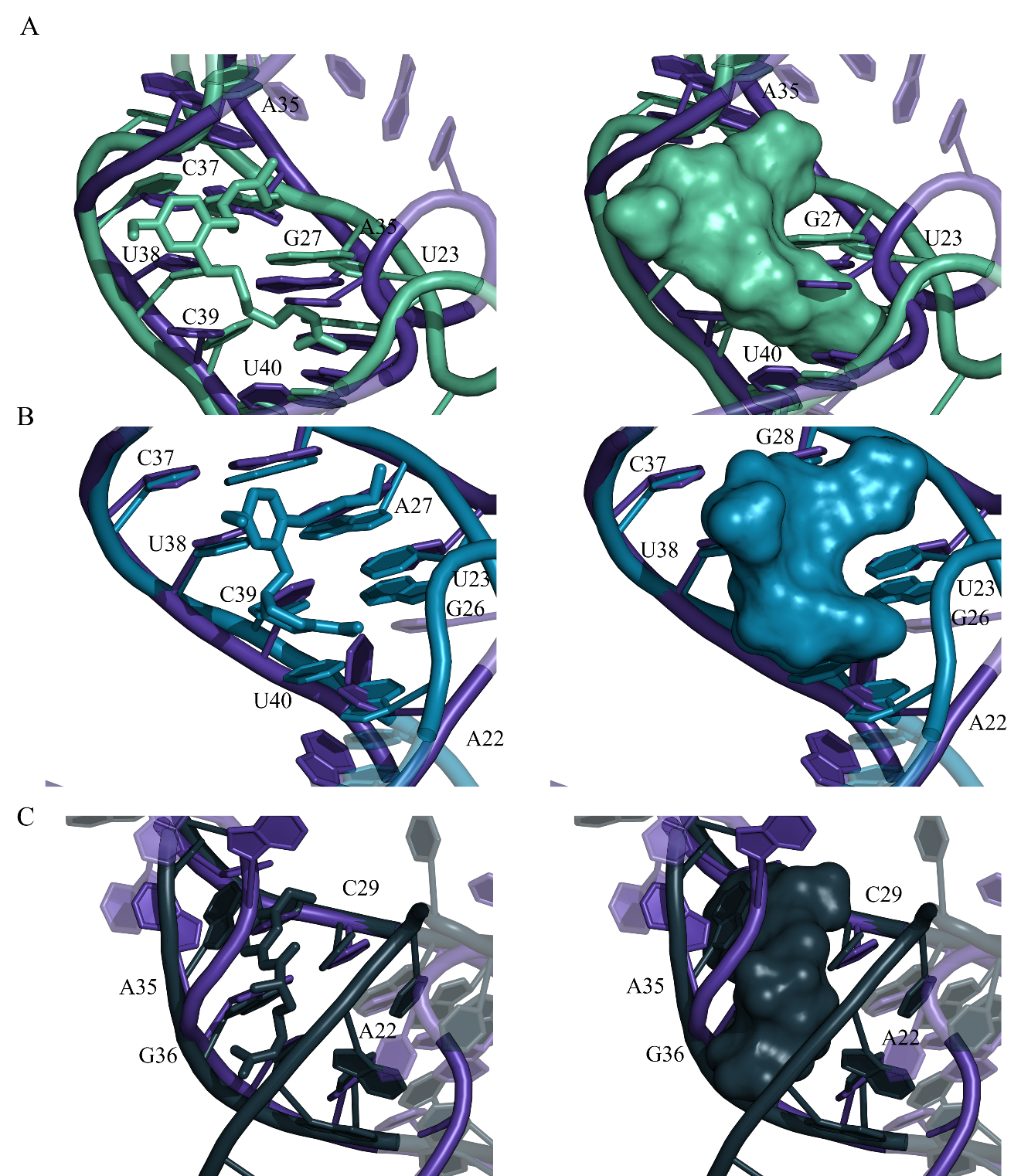

**Figure S5 Three-dimensional comparison between the closest ensemble representative and the corresponding crystal structures.** Crystal models are colored by PDB identifier. For each pocket, the left panel shows the ligand as a molecular surface to assess steric clashes and shape complementarity; the right panel shows the ligand in sticks to highlight atomic contacts. A, 1LVJ pocket; B, 1QD3 pocket; C, 1UTS pocket. Residue labels follow the original PDB numbering; the right panel shows the ligand in sticks to highlight atomic contacts. A, 1UUD pocket; B, 1UUI pocket; C, 2L8H pocket. Residue labels follow the original PDB numbering.

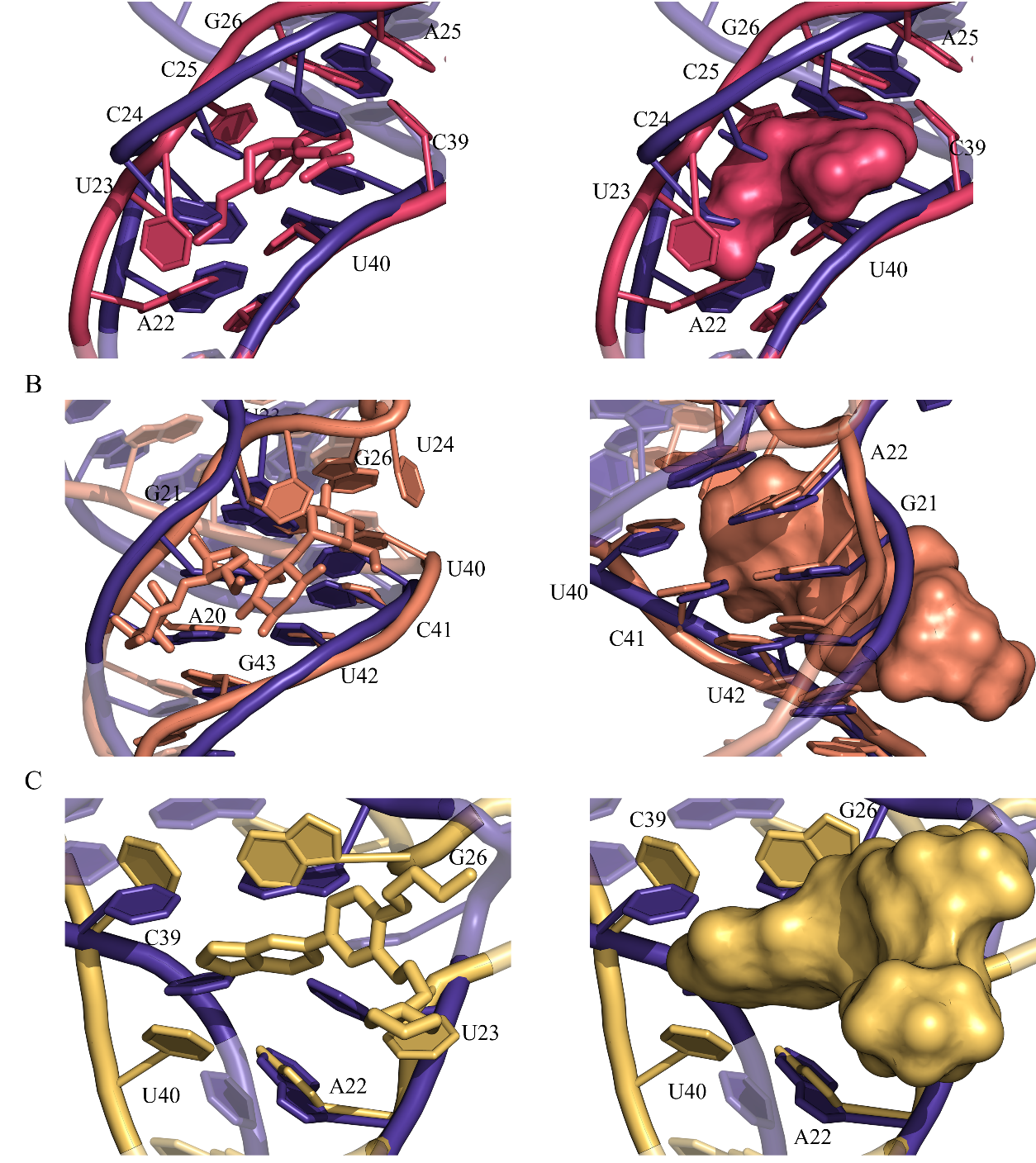

**Figure S6 Three-dimensional comparison between the closest ensemble representative and the corresponding crystal structures.** Crystal models are colored by PDB identifier. For each pocket, the left panel shows the ligand as a molecular surface to assess steric clashes and shape complementarity; the right panel shows the ligand in sticks to highlight atomic contacts. A, 1UUD pocket; B, 1UUI pocket; C, 2L8H pocket. Residue labels follow the original PDB numbering.
